# Development of conformational BRET biosensors that monitor Ezrin, Radixin and Moesin activation in real-time

**DOI:** 10.1101/2020.07.10.197707

**Authors:** Kévin Leguay, Barbara Decelle, Yu Yan He, Mireille Hogue, Hiroyuki Kobayashi, Christian Le Gouill, Michel Bouvier, Sébastien Carréno

**Affiliations:** Institute for Research in Immunology and Cancer (IRIC), Université de Montréal, P.O. Box 6128, Station Centre-Ville, Montréal, QC H3C 3J7, Canada; Cellular Mechanisms of Morphogenesis during Mitosis and Cell Motility lab; Molecular pharmacology lab; Université de Montréal, Montréal, Québec H3C 3J7, Canada; Department of Pathology and Cell Biology; Department of Biochemistry and Molecular Medicine

**Author notes:** Corresponding authors: M. Bouvier, S. Carreno. These authors contributed equally to this work (Michel Bouvier, Sébastien Carréno).

## Abstract

Ezrin, Radixin and Moesin compose the family of ERM proteins. They link actin and microtubule filaments to the plasma membrane to control signaling and cell morphogenesis. Importantly, their activity promotes invasive properties of metastatic cells from different cancer origins. Therefore, a precise understanding of how these proteins are regulated is important for the understanding of the mechanism controlling cell shape as well as providing new opportunities for the development of innovative cancer therapies. ERMs are believed to exist in two main conformational states: a close-inactive conformation in the cytoplasm that is recruited at the plasma membrane to convert into an open-active conformation. Here, we developed and characterized novel BRET-based conformational biosensors, compatible with high throughput screening, that monitor Ezrin, Radixin or Moesin individual activation in living cells. We showed that these biosensors faithfully monitor ERM activation and can be used to quantify the impact of small molecules, mutation of regulatory amino acids or depletion of upstream regulators on their activity. The use of these biosensors allowed us to uncover a novel aspect of ERM activation process involving a pool of closed-inactive ERMs stably associated with the plasma membrane. Upon stimulation, this pool serves as a cortical reserve that is rapidly activated before the recruitment of cytoplasmic ERMs.

## INTRODUCTION

Ezrin, Radixin and Moesin (ERM) are cross-linkers between the plasma membrane and the cytoskeleton. This family of proteins regulate cell morphogenesis by linking actin filaments (F-actin) and microtubules to the plasma membrane^1-3^. Thereby, ERMs regulate several important functions such as T cell activation, cell migration, neurogenesis, epithelial maintenance and cell division^1^. In a pathological context, overexpression and activation of ERM proteins participate to cancer progression^4^. While the role of ERM proteins in this process is not totally understood, their experimental inactivation was shown to reduce metastasis of cells from different cancer origin^5,6^. Therefore, blocking ERM activation or regulators acting upstream represents a promising new therapeutic avenue against cancer metastasis.

ERMs cycle between a closed-inactive form in the cytosol and an open-active form at the plasma membrane^7^. They harbor three conserved domains^1^: (i) A FERM (Band 4.1 Ezrin Radixin Moesin) domain at the amino-terminus that localizes ERMs at the plasma membrane by binding to phosphatidylinositol-(4,5)-bisphosphate [PtdIns(4,5)P_2_]^8-10^ and to membrane proteins^11,12^. This domain also binds to microtubules^3^. (ii) A central alpha-helix domain acting as a hinge region that allows switching between closed and open conformations^13^. (iii) A C-ERMAD (C-terminal Ezrin Radixin Moesin association domain) that interacts with F-actin^14^ and harbors a conserved regulatory threonine that is phosphorylated to sustain ERM opening (T567, T564 and T558 in Ezrin, Radixin and Moesin, respectively). Disruption of the interaction between the FERM and C-ERMAD opens the molecule and unmasks the actin-binding and microtubule-binding sites on C-ERMAD and FERM domains respectively.

How this activation is achieved is still not totally understood. A first model proposes that PtdIns(4,5)P_2_ recruits closed-inactive ERMs at the cortex to promote their opening by loosening the interaction between the FERM and C-ERMAD^15^. This open-active conformation is not stable and ERMs re-localize into the cytosol in their closed-inactive conformation^16^. A kinase stabilizes the open conformation at the plasma membrane by phosphorylating the conserved threonine in the C-ERMAD. Recent work has proposed an alternative model for the activation of Ezrin in cell microvilli^17^. In this model, PtdIns(4,5)P_2_ binding to Ezrin primes its activation and LOK, a Ser/Thr kinase, promotes its full opening by wedging in between the FERM domain and the C-ERMAD to physically distance these domains. Then, LOK stabilizes this open-active conformation by phosphorylating the regulatory threonine. This latter model implies that ERMs exist under a closed conformation at the plasma membrane while in the first model, ERMs are recruited and opened simultaneously.

Presently, available tools do not allow to precisely dissect the sequence of events linking the translocation to the plasma membrane and the opening and activation of ERMs. Up to now, only the last step of ERM activation is assessed using an anti-phospho antibody that targets the phosphorylated regulatory threonine. Since the surrounding region of this threonine is 100% identical in every ERMs, this antibody is not specific to the activation state of individual ERMs. Here, we aimed to develop new tools to monitor ERM activation. We elected to develop ERM biosensors based on the principle of bioluminescence resonance energy transfer (BRET)^18^ and enhanced-bystander BRET (ebBRET)^19^, since such sensors are easy to use in living cells and are compatible with high-throughput screening^20-23^. This could ultimately allow the identification of novel small compounds blocking the metastatic activity of ERMs.

We developed a collection of BRET-based biosensors that monitor the conformation and activation of individual ERM (Ezrin, Radixin and Moesin) in living cells and validated their use for high-throughput screening. Finally using these biosensors, we discovered an intermediate step in the mechanism of activation of ERM proteins. We found that a pool of closed-inactive ERMs stably associates with the plasma membrane. Upon stimulation, this intermediate pool allows a fast and local activation of ERMs that precedes the cortical recruitment and activation of cytoplasmic ERMs.

## RESULTS

### Development of enhanced-bystander BRET (ebBRET)-based biosensors to monitor the regulation of individual ERMs

As a proxy of ERM activation, we aimed to design BRET-based biosensors that can quantify Ezrin, Radixin or Moesin recruitment at the plasma membrane. To do this, we constructed the first component of the ebBRET biosensors by fusing the bioluminescent donor *Renilla* luciferase (rLucII) at the C-terminus of individual ERM (thereafter referred to as individual E,R,M-rLucII). The other component consists of the BRET fluorescent acceptor *Renilla* GFP (rGFP) targeted to the plasma membrane by the prenylation CAAX box of KRAS (rGFP-CAAX^24^) (Fig 1A). Thus, when ERM are recruited at the plasma membrane, we expected a rise in BRET signal resulting from an increased proximity between the individual Ezrin-, Radixin-, Moesin-rLucII donors (E,R,M-rLucII) and rGFP-CAAX acceptor.

**Figure 1:**
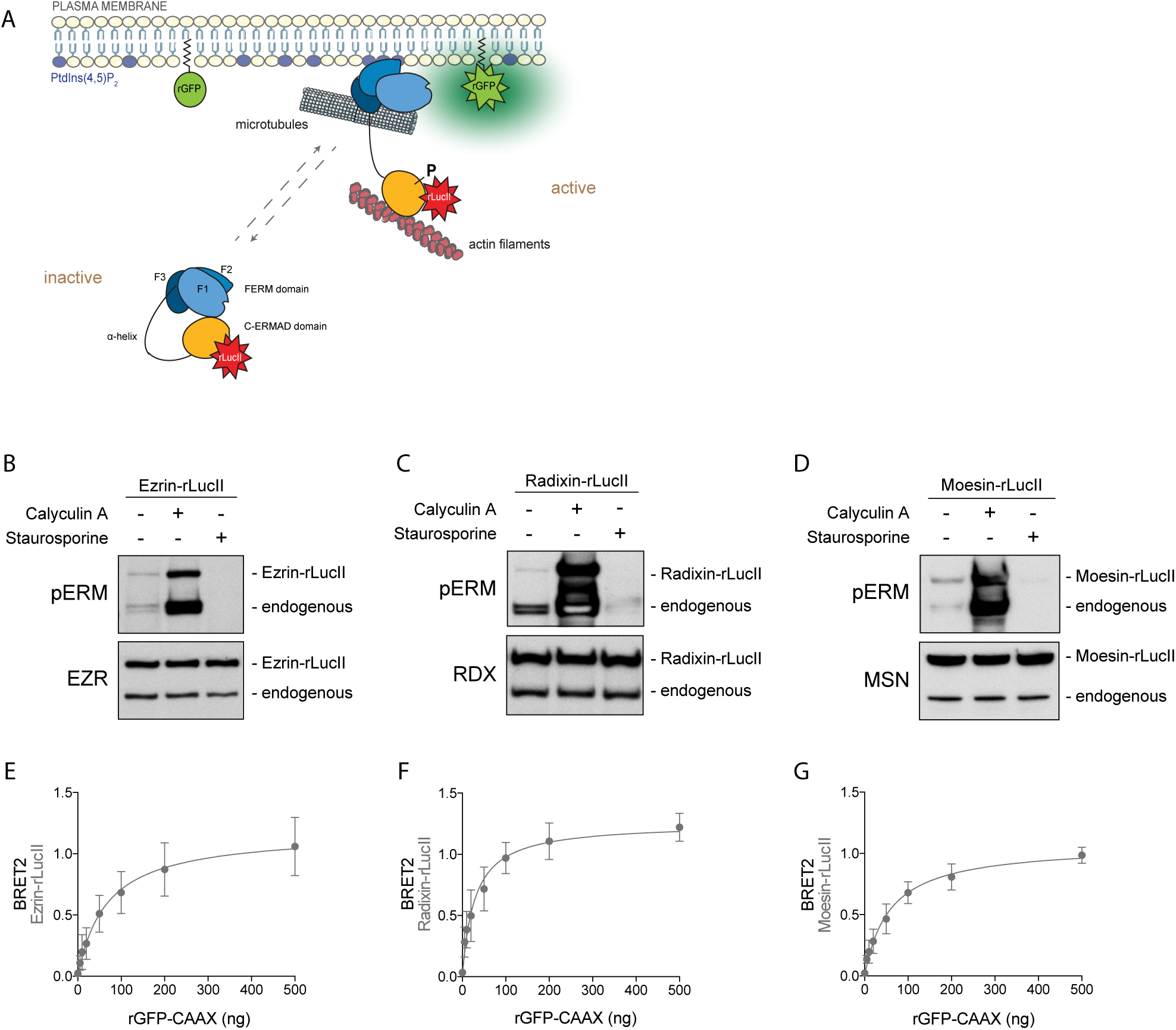
Development of BRET bimolecular biosensors to monitor the activation of Ezrin, Radixin and Moesin. (A) Illustration of ebBRET-based ERM biosensors. rLucII (donor) is fused to Ezrin, Radixin or Moesin C-terminus whereas rGFP (acceptor) is anchored at the plasma membrane with a CAAX box. (B-D) Immunoblots of HEK293T cells transfected with the indicated E,R,M-rLucII biosensors and treated with the indicated inhibitors. Immunoblots are representative of three independent experiments. (E-G) HEK293T cells were transfected with Ezrin-rLucII (E), Radixin-rLucII (F) or Moesin-rLucII (G) and the indicated amount of rGFP-CAAX. Graphs represent the mean +/- s.d. of three independent experiments.

To validate that the fusion of rLucII at E,R,M C-terminus did not affect their regulation, their phosphorylation state in response to Ser/Thr kinase and phosphatase inhibitors were tested in HEK293T cells. As shown in Fig 1, both calyculin A (which inhibits Ser/Thr phosphatases and increases ERM phosphorylation^25^) and staurosporine (which inhibits Ser/Thr kinases and prevents ERM phosphorylation^26^) modulated the phosphorylation of the regulatory threonine of individual E,R,M-rLucII to a similar extent as endogenous ERMs (Fig 1B-D). This shows that rLucII fusion does not affect the overall regulation of the individual ERM biosensors.

The ability of the biosensors to detect the association of ERM with the plasma membrane was then characterized by measuring BRET in HEK293T cells co-expressing individual E,R,M-rLucII and rGFP-CAAX. Upon addition of the rLucII substrate, we found that the BRET signal generated by these three biosensors was saturable in titration experiments with increasing amounts of the rGFP-CAAX BRET acceptor (Fig 1E-G). This progressive increase of BRET signal following a hyperbole illustrates the specificity of the signal detected and indicates that a detectable fraction of the E,R,M-rLucII donors are interacting with the plasma membrane under basal condition. This confirms that ERMs are constitutively associated with the plasma membrane, which is consistent with previous reports and with their critical role in cortical regulation.

### Chemical activation of ERMs reveals a pool of closed-inactive ERMs at the plasma membrane

To determine if the BRET signal detected under basal condition reflects the presence of closed-inactive or open-active forms at the plasma membrane, we assessed the effect of the phosphatase inhibitor on the signal. Unexpectedly, addition of calyculin A promoted a net drop of each individual ERM biosensor BRET signal compared to the vehicle (Fig 2A-C). This decrease suggests the existence of a significant fraction of ERMs that is already localized at the plasma membrane in a closed-inactive conformation. In their closed conformation, ERMs are ∼10 nm in length while they can span up to ∼40 nm when open^27^. Given that BRET efficiency varies with the 6^th^ power of the distance and the BRET R0 is approximately 5nm, if a pool of closed ERMs stably associates with the plasma membrane, ERM opening and activation could increase by four the distance between the rLucII donor and rGFP-CAAX acceptor leading to an important decrease of BRET signal (Fig 2F) as observed upon calyculin A activation.

**Figure 2:**
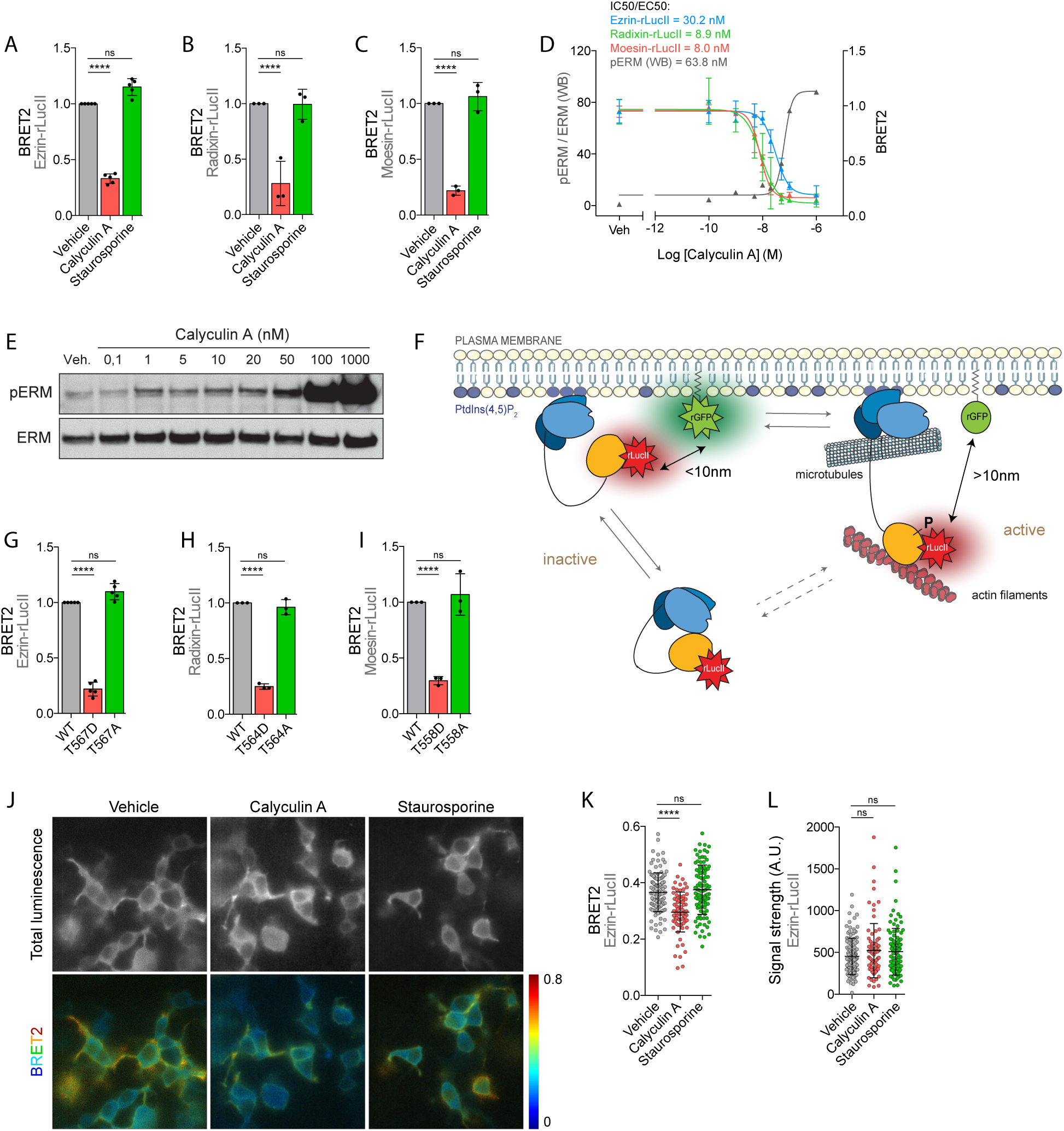
E,R,M BRET biosensors reveal a pool of inactive ERM at the plasma membrane. (A-C) E,R,M-rLucII biosensors monitor chemical activation but not chemical inhibition. HEK293T cells were co-transfected with rGFP-CAAX and Ezrin-rLucII (A), Radixin-rLucII (B) or Moesin-rLucII (C) and treated with calyculin A or staurosporine as described in Material and Methods. (D-E) BRET biosensors show similar sensitivity compared to p-ERM antibody. Cells were treated with increasing amount of calyculin A. Right axis: BRET signal measured in cells co-transfected with rGFP-CAAX and Ezrin-rLucII (blue), Radixin-rLucII (green) or Moesin-rLucII (red). Left axis: Quantification of endogenous ERM phosphorylation after immunoblotting (E). (F) Illustration of the E,R,M-rLucII BRET biosensors revealing the existence of a pool of closed ERM at the plasma membrane. (G-I) Mutation of the conserved regulatory threonine residue affects BRET signal of the ERM biosensors. HEK293T cells were co-transfected with rGFP-CAAX and a constitutively active (TD) or inactive (TA) mutant for Ezrin-rLucII (G), Radixin-rLucII (H) or Moesin-rLucII (I). (J-L) BRET-based imaging of HEK293T cells co-transfected with rGFP-CAAX and Ezrin-rLucII and treated with either calyculin A or staurosporine. BRET signal (J, bottom) and total luminescence (J, top) are quantified at the plasma membrane (K and L respectively, n>77). BRET data represent the mean +/- s.d. of multiple independent experiments. Dots represent independent experiments (A-C, G-I) or individual cells (K-L). Western blot and pERM quantification are representative of three independent experiments. P values were calculated using one-way ANOVA, Tukey’s multiple comparisons test with a single pooled variance. ****, P < 0.0001. ns, not significant.

Concentration response curves of phosphatase inhibitor revealed that the respective IC50/EC50 of calyculin A assessed by BRET and by Western blot detecting the phosphorylated regulatory threonine of ERMs are very similar [IC50 (BRET): E = 30.2 nM, R = 8.9 nM, M = 8.0 nM; EC50 (WB) = 63.8nM)] (Fig 2D-E). Such correlation with the phosphorylation event indicates that the sensors readily detect an activation process, thereby demonstrating the sensitivity of the ERM BRET biosensors. In addition, we found that individual ERM biosensors presented similar IC50 for activation by calyculin A as calculated by BRET (Fig 2D), consistent with the notion that the three ERMs have similar sensitivity to phosphatases.

To further validate that the decrease of BRET signal reveals the existence of a pool of closed-inactive ERMs at the plasma membrane, we next introduced a phospho-mimetic mutation (T/D) on the regulatory threonine of individual E,R,M-rLucII. This mutation promotes ERM opening and open-active conformation in cells^7^. Confirming our hypothesis, we found that individual ERM opening via the T/D mutation promoted a net decrease in BRET signal when compared to their wild-type counterparts (Fig 2G-I). To visualize the pool of closed-inactive ERMs at the plasma membrane, we next monitored BRET signal using BRET-based imaging^28^. As reported for endogenous Ezrin^1^, E-rLucII localized both in the cytoplasm and at the plasma membrane (Fig 2J, luminescence). As expected from the spectrometric BRET signal detected, BRET signal can readily be detected at the plasma membrane further demonstrating that rLucII fused to the C-terminus of Ezrin and rGFP-CAAX are in close proximity at the plasma membrane (Fig 2J, BRET). Also consistent with the spectrometric data presented above, upon calyculin A activation, Ezrin biosensor BRET signal decreased specifically at the plasma membrane while E-rLucII signals remained constant (Fig 2J-L). This indicates that Ezrin did not leave the plasma membrane but rather transitioned into an open conformation upon phosphatase inhibition. Altogether, these findings demonstrate that ERMs stably associate with the plasma membrane in their closed conformation.

We next assessed if these ERM biosensors could also monitor ERM inhibition. According to our finding that a pool of inactive ERMs associates with the plasma membrane, an increase of BRET signal should follow ERM inhibition since the closing of ERMs at the plasma membrane will bring the luciferase donor closer to the rGFP acceptor. While staurosporine promotes ERM dephosphorylation and inactivation (Fig 1B-D), we did not detect any significant changes in the BRET signal after staurosporine treatment with any of the individual E,R,M-rLucII biosensors (Fig 2A-C). BRET-based imaging confirmed these observations since we did not measure any significant changes in E-rLucII biosensor BRET signal at the plasma membrane upon staurosporine treatment (Fig 2J-K) nor any change in its distribution (Fig 2L). In addition, the introduction of a non-phosphorylatable mutation of the regulatory threonine (T/A) in the individual E,R,M-rLucII constructs inactivating ERMs^7^ did not significantly affect the BRET signal of each bimolecular sensors in comparison with their wild-type counterparts (Fig 2G-I). This lack of signal variation following inhibition could result from an equilibrium between an increase of signal from the plasma membrane associated-inactive conformation and a decrease of signal due to the detachment of ERMs from the plasma membrane to the cytoplasm.

### Engineering ebBRET-based biosensors to monitor ERM closing at the plasma membrane

The equilibrium between inactive ERMs at the plasma membrane and in the cytoplasm makes difficult to measure the closing of ERMs at the plasma membrane using the first ebBRET-based biosensors. To monitor this closing, we designed a second set of ebBRET-based ERM biosensors that are restricted to the plasma membrane. To this aim, we stably associated the individual E,R,M-rLucII fusions to the plasma membrane using a myristoylation (Myr) motif followed by a polybasic (PB) amino acid sequence (thereafter referred to as individual MyrPB-E,R,M-rLucII) (Fig 3A). With these new biosensors, we expect to measure an increase in BRET signal upon ERM inactivation and a decrease upon activation. In control cells, we found that the BRET signal of these three biosensors was saturable in titration experiments showing the specificity of the BRET signal (Fig 3B-D). According to the prediction, ERM inactivation by staurosporine significantly increased BRET signal of individual MyrPB-E,R,M-rLucII sensors (Fig 3B-D). To verify that this effect was not due to an unexpected modification of the composition of the plasma membrane caused by staurosporine treatment, we constructed a control biosensor composed of rLucII directly fused to the MyrPB plasma membrane targeting motif (MyrPB-rLucII). Treatment with staurosporine had no effect on the BRET signal between this plasma membrane-targeted Rluc and rGFP-CAXX (Fig 3E). As observed with the first ebBRET-based biosensors (Fig 2A-C), ERM activation with calyculin A decreased BRET signal of the three MyrPB-E,R,M-rLucII biosensors (Fig 3B-D) while it did not affect the BRET signal of the control MyrPB-rLucII biosensor (Fig 3E). We then assessed the respective EC50/IC50 of staurosporine and calyculin A on individual MyrPB-E,R,M-rLucII and endogenous ERMs measured by BRET and Western blot respectively. We found that the endogenous ERMs and MyrPB-E,R,M-rLucII biosensors presented similar EC50/IC50 for calyculin A and staurosoporine within the nM range (Fig 3F-H). Altogether, these results suggest that this second set of ebBRET-based biosensors monitor both activation and inhibition following treatment with phosphatase and kinase inhibitors, respectively.

**Figure 3:**
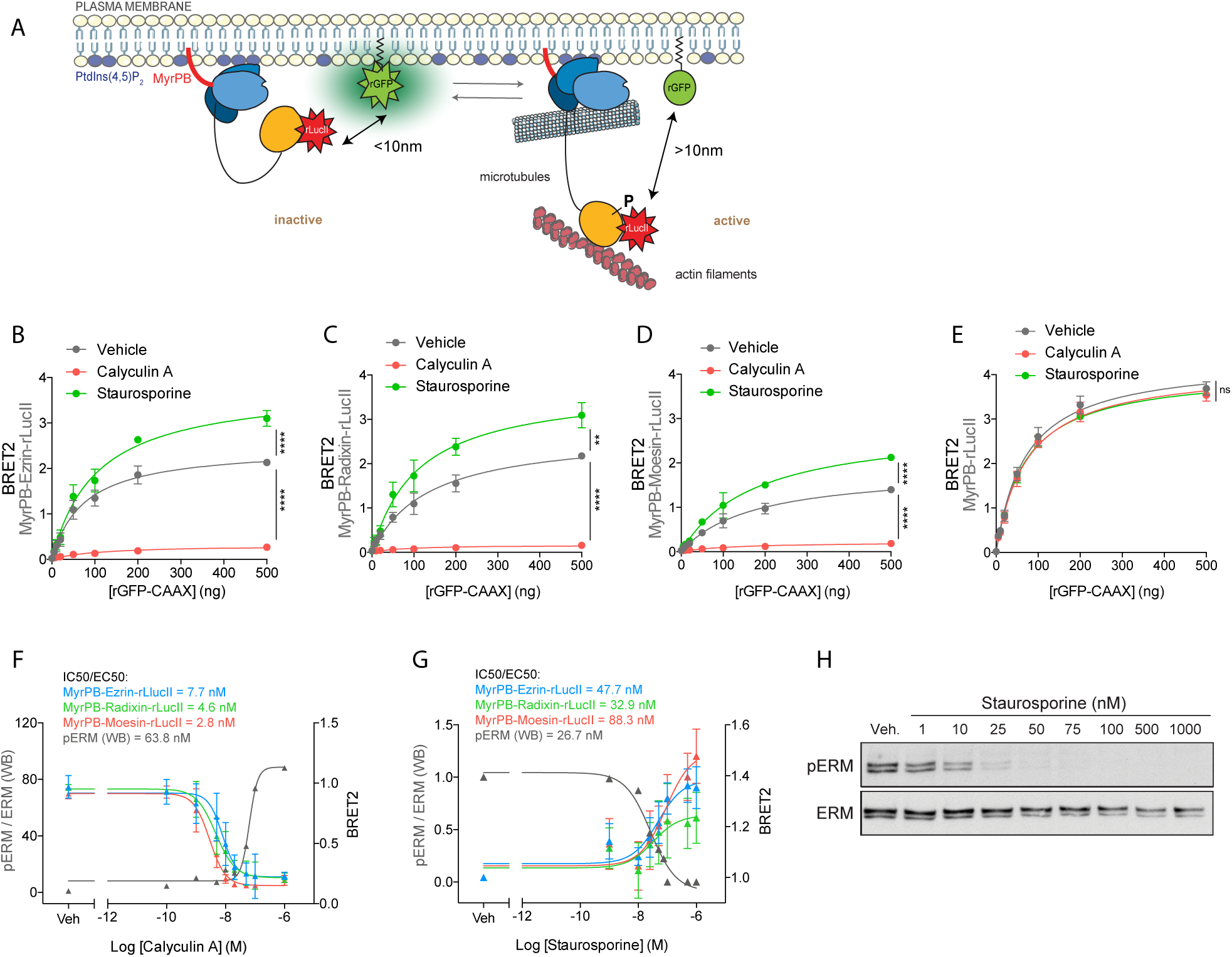
MyrPB-E,R,M-rLucII biosensors monitor both ERM activation and inhibition. (A) Illustration of the MyrPB-E,R,M-rLucII BRET biosensors. E,R,M-rLucII constructs are anchored to the plasma membrane with a myristoylation motif followed by a polybasic amino acid sequence. This anchorage avoids the shuttling of E,R,M-rLucII between the plasma membrane and the cytosol and allows the monitoring of conformational changes at the plasma membrane. (B-E) MyrPB-E,R,M-rLucII biosensors respond specifically to both activation and inhibition. HEK293T cells were co-transfected with MyrPB-Ezrin-rLucII (B), MyrPB-Radixin-rLucII (C), MyrPB-Moesin-rLucII (D) or MyrPB-rLucII (E) and indicated amount of rGFP-CAAX (acceptor) and treated with calyculin A or staurosporine. (F-H) Plasma membrane-anchored ERM BRET biosensors show similar sensitivity compared to p-ERM antibody upon chemical activation and inhibition. Cells were treated with increasing amount of calyculin A (F) or staurosporine (G) before BRET reading. (F-G) Right axis: BRET signal measured in cells co-transfected with rGFP-CAAX and MyrPB-Ezrin-rLucII (blue), MyrPB-Radixin-rLucII (green) or MyrPB-Moesin-rLucII (red). Left axis: Quantification of endogenous ERM phosphorylation after immunoblotting (H and Fig 2E). BRET signal curves represent the mean +/- s.d. of three independent experiments. Western blot and p-ERM quantification are representative of three independent experiments. P values were calculated using one-way ANOVA, Tukey’s multiple comparisons test with a single pooled variance. **, P < 0.01. ****, P < 0.0001. ns, not significant.

### The MyrPB-Ezrin-rLucII biosensor probes Ezrin conformational changes at the plasma membrane

Ezrin being the most characterized ERM during cancer metastasis^4^, we decided to focus on this paralogue to further our understanding of its regulation. We tested if the MyrPB-Ezrin-rLucII BRET signal can be modulated by introducing T567D phospho-mimetic or T567A non-phosphorylatable mutations to change its conformation. Confirming that the MyrPB-Ezrin-rLucII biosensor probes Ezrin conformational states, we observed that BRET signal decreased for the Ezrin^T567D^ mutant and increased for the Ezrin^T567A^ biosensor mutant (Fig 4A). As expected, we also found that these Ezrin phospho-mutants were no longer sensitive to calyculin A or staurosporine treatments (Fig S1).

**Figure 4:**
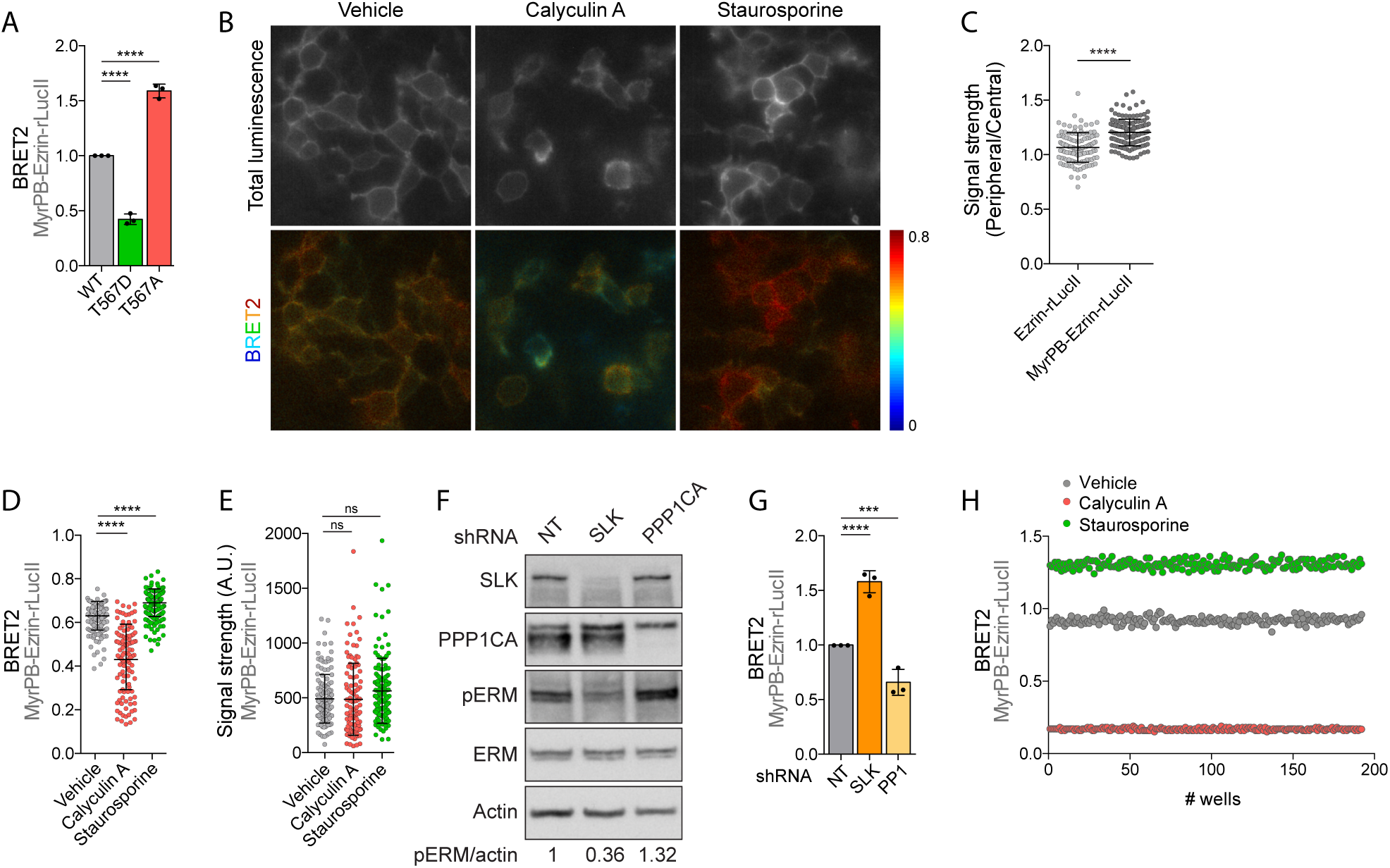
MyrPB-Ezrin-rLucII biosensor probes Ezrin conformational changes at the plasma membrane. (A) MyrPB-Ezrin-rLucII biosensor is sensitive to mutation of its regulatory conserved threonine residue. HEK293T cells were co-transfected with rGFP-CAAX and a constitutively active (T567D) or inactive (T567A) mutant of MyrPB-Ezrin-rLucII. (B-E) BRET-based imaging of HEK293T cells co-transfected with rGFP-CAAX and MyrPB-Ezrin-rLucII and treated with either calyculin A or staurosporine. BRET signal (B, bottom) and total luminescence (B, top) are quantified at the plasma membrane (D and E respectively, n>110). (C) Enrichment of rLucII activity at the plasma membrane over the cytoplasm is quantified in cells transfected with either Ezrin-rLucII (see Fig 2J) or MyrPB-Ezrin-rLucII. (F-G) Depletion of ERM known regulators affects both their phosphorylation and MyrPB-Ezrin-rLucII opening measured by BRET. HEK293T cells stably expressing shRNAs directed against indicated proteins were subjected to immunoblotting (F) and BRET measurement after co-transfection with rGFP-CAAX and MyrPB-Ezrin-rLucII (G). (H) Determination of Z’ score of MyrPB-Ezrin-rLucII and rGFP-CAAX bimolecular biosensor stably co-transfected in HEK293T cells and treated with calyculin A or staurosporine in 384-well plates. Graphs represent the mean +/- s.d. of three independent experiments. Dots represent independent experiments (A, G), individual cells (C-E) or individual wells (H). P values were calculated using one-way ANOVA, Tukey’s multiple comparisons test with a single pooled variance. ***, P < 0.001. ****, P < 0.0001. ns, not significant.

Using BRET-based imaging, we confirmed that MyrPB-Ezrin-rLucII is significantly enriched at the plasma membrane (Fig 4B-C) when compared with the Ezrin-rLucII biosensor (Fig 2J). Furthermore, MyrPB-Ezrin-rLucII BRET signal at the plasma membrane significantly decreased upon calyculin A activation and increased upon staurosporine inactivation, without any significant change of the signal luminescence intensity of MyrPB-Ezrin-rLucII (Fig 4B, D-E).

Having shown that the MyrPB-Ezrin-rLucII biosensor faithfully monitor Ezrin conformation changes upon chemical activation or inhibition, we then tested if the BRET signal of this biosensor can be modulated by depleting the upstream regulators of ERMs. We previously reported that the regulatory threonine of dMoesin, the sole ERM protein in Drosophila, is phosphorylated by the Ser/Thr kinase Slik and is dephosphorylated by the Ser/Thr phosphatase PP1-87B^10,29^. In mammals, SLK, the human orthologue of Slik and the close paralogue of LOK, phosphorylates ERM proteins^30^ whereas PPP1CA, an orthologue of PP1-87B, dephosphorylates them^31^. We thus knocked-down SLK and PPP1CA by shRNA and we confirmed that these orthologues also regulate ERM phosphorylation in HEK293T cells by Western-blot (Fig 4F). Confirming the robustness of the individual MyrPB-E,R,M-rLucII biosensors, we observed that the BRET signal associated to Ezrin, Radixin and Moesin increased upon SLK depletion and decreased upon PPP1CA depletion (Fig 4G and S2). Altogether, these results show that membrane restricted ERM BRET biosensors are universal and precisely monitor both activation and inhibition of individual ERM by sensing their open and closed conformation at the plasma membrane.

Finally, we assessed if the MyrPB-Ezrin-rLucII ebBRET-based biosensor is suitable for high-throughput screening. To this aim, we measured the Z-factor^32^ (Z’) of activation and inhibition in HEK293T cells seeded in 384-well plate using automated procedures (see material and methods). Vehicle (DMSO), calyculin A or staurosporine were randomly distributed on cells stably expressing the MyrPB-Ezrin-rLucII and rGFP-CAAX bimolecular ebBRET-based biosensor. One hour after treatment, BRET signal was read and we found that calyculin A and staurosporine induced highly reproducible decrease or increase of MyrPB-Ezrin-rLucII BRET signal, respectively, when compared with vehicle alone (Fig. 4F). Both Z factor of activation (Z’=0.87 upon calyculin A treatment) and inhibition (Z’=0.59 upon staurosporine treatment) are compatible with high-throughput screening procedures showing that this biosensor will be valuable to identify compounds that modulate Ezrin activation after screening of compound libraries.

### Closed-inactive cortical Ezrin serve as a reserve of proteins that can be quickly and locally activated

The discovery of a pool of closed ERMs that stably associates with the plasma membrane opens the possibility that these ERMs serve as a reserve of proteins that can be rapidly and locally activated. We explored this hypothesis by comparing the timing of Ezrin conformational opening by BRET with the timing of Ezrin recruitment at the plasma membrane using live-cell imaging in HEK293T cells expressing Ezrin-GFP. Upon phosphatase inhibition, cortical closed-inactive Ezrin opened very rapidly as the BRET signal started to decrease at 1 min (the earliest timepoint tested) and reached a plateau at ∼4 min for both the Ezrin-rLucII and MyrPB-Ezrin-rLucII biosensors (Fig 5A). In the meantime, Ezrin-GFP was recruited at the plasma membrane only after 9 minutes of activation (Fig 5B-C). In parallel, we observed that upon activation, phosphorylation of the regulatory threonine of ERMs also started very rapidly (∼ 2min, the earliest timepoint tested) but reached a plateau only after 10 min; 6 min after the cortical pool of closed-inactive ERMs was fully activated (10 min vs 4 min) (Fig 5A & D). Although these differences of timing could be due to the use of different molecular tools, we hypothesized that the cortical pool of plasma membrane associated-inactive Ezrin is activated before the *de novo* recruitment of cytoplasmic inactive Ezrin.

**Figure 5:**
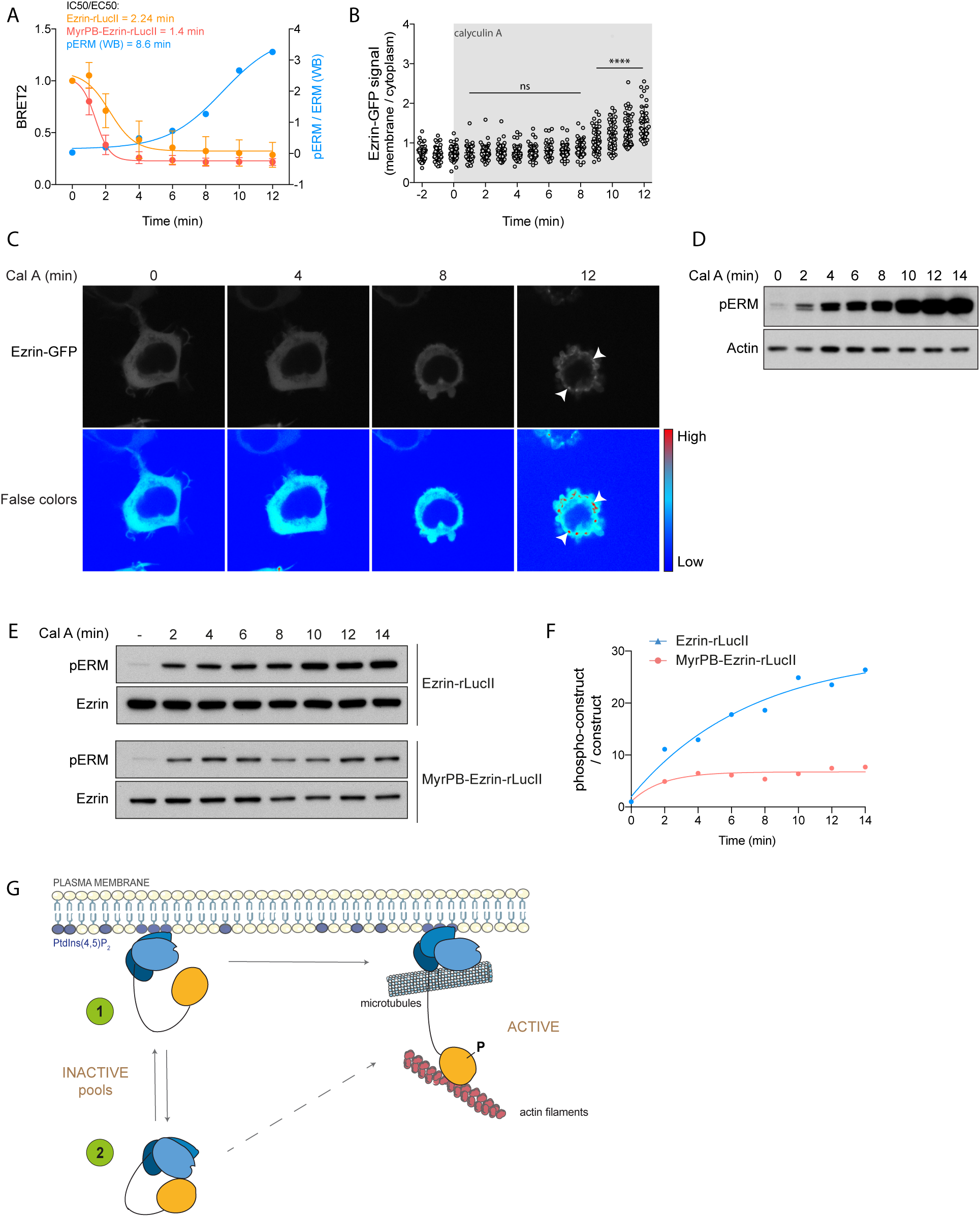
The pool of inactive ERM at the plasma membrane is activated before recruitment of cytoplasmic ERM. (A) BRET-based real-time analysis of Ezrin-rLucII (orange) and MyrPB-Ezrin-rLucII (red) activation induced by calyculin A compared to ERM phosphorylation measured by western bolt (blue, quantification of immunoblot in D). HEK293T cells were co-transfected with rGFP-CAAX and Ezrin-rLucII or MyrPB-Ezrin-rLucII, then treated with calyculin A. BRET signal curves represent the mean +/- s.d. of three independent experiments. (B-C) Time-lapse analysis of Ezrin-GFP recruitment at the plasma membrane by live-cell imaging. HEK293T cells were transfected with Ezrin-GFP and treated with calyculin A. Images (C) were acquired every minute and GFP signal intensity at the plasma membrane was quantified using Image J (B). Graph represents the mean +/- s.d. of three independent experiments. Each individual dot represents the measure for one individual cell (n>45). (D) Representative immunoblot against pERM at the specified time after addition of calyculin A. (E-F) Time-lapse analysis of Ezrin-rLucII and MyrPB-Ezrin-rLucII phosphorylation on their regulatory residue after addition of calyculin A during the indicated time. (G) Inactive-closed ERMs are distributed between the plasma membrane (1) and the cytosol (2). Upon stimulation, cortical closed-inactive ERMs (1) are first activated followed by *de novo* recruitment of inactive cytosolic ERMs (2). Immunoblots and p-ERM quantifications are representative of three independent experiments. P values were calculated using one-way ANOVA, Tukey’s multiple comparisons test with a single pooled variance. ****, P < 0.0001. ns, not significant.

We tested this hypothesis by comparing the profile of phosphorylation of Ezrin-rLucII with those of the plasma membrane associated MyrPB-Ezrin-rLucII after activation (Fig 5E-F). We found that while the maximal phosphorylation of MyrPB-Ezrin-rLucII is achieved rapidly after 2 min of activation by calyculin A, Ezrin-rLucII phosphorylation increases more gradually to reach a plateau at 10 min. Altogether, these data show that the cortical pool of closed-inactive Ezrin is activated first. Then, upon sustained activation, cytoplasmic Ezrin is recruited at the plasma membrane to be concomitantly phosphorylated and activated (Fig 5G).

## DISCUSSION

Here, we developed and characterized a set of ebBRET-based biosensors that measure the activation of individual ERMs. These biosensors locally monitor the conformational changes of ERMs at the plasma membrane that dictate their activity. We demonstrated that these biosensors faithfully measure ERM regulation in live cells following activation and inhibition. In addition, these biosensors revealed a pool of closed-inactive ERMs associated with the plasma membrane that can be rapidly and locally open and activated upon stimulation. This finding adds a supplementary layer of refinement in the model of regulation of ERM proteins.

While characterizing the first set of ERM biosensors, we discovered that a pool of ERM proteins associates with the plasma membrane under a closed conformation. A recent work established that after Ezrin recruitment at the plasma membrane of microvilli, the Ser/Thr kinase LOK opens ERM proteins by wedging in between the FERM domain and the C-ERMAD^17^. While the available molecular tools did not allow to test it, this model implies that Ezrin is already associated with the plasma membrane in its closed conformation before activation. Our data confirm this model by demonstrating that a pool of inactive-closed ERMs is stably associated with the cortex. Beyond the regulation of Ezrin in microvilli, our discovery reveals an accurate and universal model for ERM activation (see model in Fig 5G).

Generating one specific cell shape that promotes one specific biological function relies on the integration of actin filaments at the plasma membrane by proteins such as ERMs. Ezrin, Radixin and Moesin regulate several functions that require a rapid remodeling of cell shape such as cell division or cell migration. In agreement with the rapid timeline associated with these functions, we discovered that the cortical pool of closed Ezrin can be activated within 1 min (see Fig 5A), even before the other pool of cytosolic Ezrin is recruited and activated at the plasma membrane (see Fig 5B-C and model in Fig 5G). Interestingly, we observed that, upon calyculin A treatment, BRET signal associated with Ezrin-rLucII is stable overtime (see Fig 5A) even after 9min, when cytosolic Ezrin is recruited (see Fig 5B-C). Since BRET signal occurs when the rLucII and rGFP-CAAX are in close proximity at the plasma membrane, this suggests that the recruitment of cytosolic closed-inactive ERMs is immediately followed by their opening and activation. This discovery contributes to a better comprehension of how ERMs are open and activated at the plasma membrane. Yet, we still need more information to understand how the trafficking of cytosolic ERMs toward the plasma membrane can be triggered after stimulation.

Until now, the main approach used to study ERM activation was to follow the phosphorylation of the C-ERMAD regulatory threonine residue using a specific phospho-antibody by western-blot or by immunofluorescence. However, this approach based on this p-ERM antibody presents several limitations: (i) this antibody does not discriminate between the phosphorylated form of the three ERMs. Therefore, it is not possible to visualize the level of activation of each individual ERM in cells by immunofluorescence. In addition, since Ezrin and Radixin have a similar apparent molecular weight, they cannot be separated by regular SDS-PAGE. Here, we developed biosensors that monitor the activity of each ERM independently. (ii) ERM phosphorylation at their regulatory C-ERMAD threonine was shown to stabilize their active open conformation, and therefore is commonly associated with their activation state. However, ERMs were also found to be activated without being phosphorylated at this conserved residue. For instance, phosphorylation of Threonine 235 of the FERM domain of Ezrin by CDK5 was shown to be sufficient to open and activate this ERM^33^. The conformational biosensors directly monitor the closed-inactive and open-active forms of ERMs at the plasma membrane and thus bypass the requirement that the C-ERMAD threonine is phosphorylated. (iii) The use of antibodies is not compatible with real-time analysis in live cells. Thus, assessing sequential activation of ERMs by western-blot or immunofluorescence requires complex experimental approaches with different samples for every timepoints. This greatly increases the variability between conditions and experiments. In the present study, we showed that the ERM bimolecular biosensors allow monitoring of ERM activation within the same cell population over time. We took advantage of this to compare the activation kinetics of the different pools of Ezrin. In addition, using BRET-based imaging, we showed that the level of ERM activation can be visualized in real-time in living cells.

Last, ERMs were recently identified as promising therapeutic targets against metastasis^4,6,34^. The development of new ERM therapeutic inhibitors requires high-throughput screening procedures that are not compatible with the existing experimental approaches to study ERMs. BRET-based biosensors have already been widely used in the last years in genetic and chemical screens. Here, we developed ERM bimolecular biosensors that present high and reproducible BRET signal in 384-well plates, compatible with high-throughput screening. We are currently using these new BRET-based biosensors to identify new compounds targeting ERM activity and therefore their potential ability to inhibit the metastatic progression associated with ERMs.

## ACKNOWLEDGMENT

This work has been supported by CCSRI Innovation Grant (705892 to S.C.) and a Foundation Grant from the Canadian Institute for Health Research (148431 to M.B). K.L held a doctoral scholarship from IRIC and from Montreal University’s Molecular Biology Program as well as a ESP fellowship from Montreal University. Y.Y.H held a master scholarship from IRIC. MB holds the Canada Research Chair in Signal Transduction and Molecular Pharmacology.

## AUTHOR CONTRIBUTIONS

M.B and S.C. managed the project. K.L., B.D., C.L.G, M.B. and S.C. conceptualized and designed the experiments. K.L., B.D., Y.Y.H, M.H., H.K. and C.L.G performed the experiments. K.L, B.D., C.L.G, M.B. and S.C. analyzed the data. K.L, B.D. and S.C. prepared the figures for the manuscript. K.L, M.B. and S.C. wrote the manuscript with input from all coauthors.

## MATERIAL & METHODS

### Reagents and inhibitors

Coelenterazine 400a (Deep Blue C) and methoxy e-CTZ (Prolume Purple) were purchased from NanoLight Technology (#340 and #369, respectively). Calyculin A was purchased from Sigma (#C5552). Staurosporine was purchased from ApexBio (#A8192).

### DNA constructs

The rGFP-CAAX construct was previously described^24^. Ezrin-rLucII, Radixin-rLucII and Moesin-rLucII constructs were obtained by subcloning PCR amplified Ezrin, Radixin or Moesin into pCDNA 3.1 hygro(+) GFP10-rLucII vector^35^ digested with NheI and AgeI to replace GFP10. MyrPB-E,R,M-rLucII constructs were obtained by adding a myristoylation sequence followed by a polybasic motif (MyrPB: MGCTLSAEDKAAVERSKMAVQSPKKGLLQRLFKRQHQTIPRVAVQNAAIRSGGSGGSGGSGGSNAAIRS) to the E,R,M-rLucII constructs. The negative control MyrPB-rLucII was obtained by subcloning the MyrPB sequence into pCDNA 3.1 hygro(+) GFP10-rLucII vector. The constitutively active and inactive (MyrPB-)Ezrin-rLucII mutants (MyrPB-)Ezrin^*T567D*^-rLucII and (MyrPB-)Ezrin^T567A^-rLucII were obtained by inverse PCR from the wildtype constructs. Primers used were: (forward) 5’-AAGATCTGCGGCAGATCCGGCAGGGCAACACCAAGC-3’ and (reverse) 5’-TGTACTTGTCCCGGCCTTGCCTCATGTTCTCG-3’ for T567D mutant; (forward) 5’-AAGCTTTGCGGCAGATCCGGCAGGGCAACACCAAGC-3’ and (reverse) 5’-TGTACTTGTCCCGGCCTTGCCTCATGTTCTCG-3’ for T567A mutant. For Radixin^T564D^-rLucII and Radixin^T564A^-rLucII, primers used were: (forward) 5’-AAGATCTGCGACAGATTCGACAAGGCAATACAAAGC-3’ and (reverse) 5’-TGTACTTATCACGGCCTGCTTTAACATTCTCAGC-3’ for T564D mutant; (forward) 5’-AAGCTTTGCGACAGATTCGACAAGGCAATACAAAGC-3’ and (reverse) 5’-TGTACTTATCACGGCCTGCTTTAACATTCTCAGC-3’ for T564A mutant. Finally, primers used for Moesin^T558D^-rLucII and Moesin^T558A^-rLucII mutants were: (forward) 5’-AAGATCTGCGCCAGATCCGGCAGGGCAACACCAAGC-3’ and (reverse) 5’-TGTATTTGTCTCGGCCCAGTCGCATGTTCTCAGC-3’ for T558D mutant; (forward) 5’-AAGCTTTGCGCCAGATCCGGCAGGGCAACACCAAGC-3’ and (reverse) 5’-TGTATTTGTCTCGGCCCAGTCGCATGTTCTCAGC-3’ for T558A mutant. Ezrin-GFP was obtained by subcloning PCR amplified Ezrin into pEGFP-N1 vector. All PCR reactions were performed using Phusion High-Fidelity DNA Polymerase (New England Biolabs). MISSION shRNA constructs were obtained from Sigma in pLKO.1-puro vectors: SLK (TRCN0000000897), PPP1CA (TRCN0000002453).

### Cell culture, transfection and infection

HEK293T human kidney cells were maintained in Dulbecco’s modified Eagle’s medium (DMEM; 4.5g/L D-Glucose, L-Glutamine, 110mg/L sodium pyruvate; Thermofisher #11995073) supplemented with 10% fetal bovine serum (FBS; Life Invitrogen #12483020) and 1% penicillin-streptomycin antibiotics (ThermoFisher #15070063) at 37°C with 5% CO_2_. For transfection, cells (350,000 cells/mL) were resuspended in DMEM supplemented with 10% FBS and transfected with 1µg of total DNA (50ng rLucII construct, 300ng rGFP-CAAX and 650ng salmon sperm DNA) using linear polyethyleneimine (PEI, Alfa Aesar #43896) as transfecting agent with a PEI:DNA ratio of 3:1. Cells were then plated in white 96-well culture plates (VWR #82050-736) at a concentration of 35,000 cells per well and were incubated for 48h prior BRET measurement.

For lentiviral infection, cells were seeded at 5,000 cells per well in 96-well culture plates in DMEM supplemented with 10% FBS and 5µg/ml polybrene (Sigma #H9268). Lentiviruses were added to the medium and cells were incubated for 24h. The next day, cells were transfected with 100ng total DNA using PEI (as described earlier) and incubated for an additional 24h. Finally, infected cells were selected with 2µg/ml puromycin (EMD Millipore #540222) for 48h.

### BRET measurements

Transfected cells with BRET biosensors were washed with Hank’s balanced salt solution (HBSS, ThermoFisher #14065056). Calyculin A and staurosporine were added at 100nM for 10min and 30min respectively, unless mentioned otherwise. Coelenterazine 400a diluted in HBSS was added 5min prior the reading at a final concentration of 2.5µM. BRET was monitored with a Tecan Infinite 200 PRO multifunctional microplate reader (Tecan) equipped with BLUE1 (370-480nm; donor) and GREEN1 (520-570nm; acceptor) filters. BRET signal was calculated as a ratio by dividing the acceptor emission over the donor emission. For automated procedures, inhibitors were added using an Echo 555 acoustic dispenser (Labcyte) while BRET2 was monitored with a Synergy NEO HTS microplate reader (Biotek).

### BRET-based imaging

Forty-eight hours before imaging, HEK293T cells were transfected as described before with rLucII-tagged BRET donors and rGFP-CAAX acceptor and plated in 35mm glass bottom culture dishes (MatTek #P35GC-0-14-C). The day of the experiment, cells were washed with HBSS and inhibitors were added. rLucII substrate methoxy e-CTZ was added at a final concentration of 10µM prior the acquisition. Images and analysis were obtained as previously described^28^.

### Immunoblotting

After treatment, HEK293T cells were washed with ice-cold phosphate-buffered saline (PBS) and lysed in TLB buffer (40 mM HEPES, 1 mM EDTA, 120 mM NaCl, 10 mM NaPPi, 10% glycerol, 1% TritonX-100, 0.1% SDS) supplemented with both phosphatase and protease inhibitors (phosphatase inhibitor cocktail (PIC, Sigma #P2850), 1 mM sodium orthovanadate (Na_3_VO_4_, Sigma #S6508), 5 mM β-Glycerophosphate (Sigma #G6251), 1 mM phenylmethylsulfonyl fluoride (PMSF, Sigma #P7626) and anti-protease cocktail (Sigma #4693132001)). After Bradford quantification, cell lysates were diluted with sample buffer (200mM TrisHCl 1M pH6.8, 8% SDS, 0.4% bromophenol blue, 40% glycerol and 412mM β-mercaptoethanol). Samples were resolved by 8% SDS-PAGE and transferred to nitrocellulose membranes (pore 0.2μm, VWR #27376-991). Membranes were blocked in TBS-Tween (25mM Tris-HCl pH 8, 125mM NaCl, 0.1% Tween 20) supplemented with 2% BSA for one hour before overnight incubation with primary antibodies at 4°C. Primary antibodies used are: rabbit anti-ERM (1:1000, Cell Signaling #3142), rabbit anti-Ezrin (1:1000, Cell Signaling #3145), rabbit anti-Radixin (1:1000, Cell Signaling #2636), rabbit anti-Moesin (1:1000, Cell Signaling #3150), rabbit anti-phospho-ERM (1:5000,^10^), mouse anti-Actin (1:5000, Sigma #MAB1501), rabbit anti-SLK (1:500, Cerdalane #A300-499A), rabbit anti-PPP1CA (1:1000, Cedarlane #A300-904A-M). Goat anti-rabbit HRP antibody (1:10000, Santacruz #sc-2004) and goat anti-mouse HRP (1:10000, Santacruz #sc-516102) were used as secondary antibodies. Finally, protein detection was performed using Amersham ECL Western Blotting detection reagent (GE Healthcare #CA95038-564L). Immunoblot quantifications were performed using ImageJ software (NIH).

### Live imaging

Forty-eight hours before the experiment, HEK293T cells were transfected with Ezrin-GFP in 35mm glass bottom culture dishes (MatTek #P35GC-0-14-C) as described before. The day of the experiment, images were acquired every minute using Yokogawa CSU-X1 5000 spinning disk confocal microscope. Quantification of the signal at the plasma membrane was performed using Image J software (NIH).

### Data analysis

All data from BRET and western blot analysis were analyzed using GraphPad PRISM software (GraphPad Software, La Jolla, CA, USA). All data are represented by the mean +/- s.d. of multiple independent experiments. Microscopy images were prepared using Image J software (NIH) and Photoshop (Adobe).

## REFERENCES

1 Fehon, R. G., McClatchey, A. I. & Bretscher, A. Organizing the cell cortex: the role of ERM proteins. Nat Rev Mol Cell Biol 11, 276–287, doi:nrm2866 [pii] 10.1038/nrm2866 (2010).

2 Polesello, C. & Payre, F. Small is beautiful: what flies tell us about ERM protein function in development. Trends Cell Biol 14, 294–302, doi: 10.1016/j.tcb.2004.04.003 (2004).

3 Solinet, S. et al. The actin-binding ERM protein Moesin binds to and stabilizes microtubules at the cell cortex. J Cell Biol 202, 251–260, doi: 10.1083/jcb.201304052 (2013).

4 Clucas, J. & Valderrama, F. ERM proteins in cancer progression. J Cell Sci 127, 267–275, doi: 10.1242/jcs.133108 (2014).

5 Bulut, G. et al. Small molecule inhibitors of ezrin inhibit the invasive phenotype of osteosarcoma cells. Oncogene 31, 269–281, doi: 10.1038/onc.2011.245 (2012).

6 Ghaffari, A. et al. Intravital imaging reveals systemic ezrin inhibition impedes cancer cell migration and lymph node metastasis in breast cancer. Breast cancer research : BCR 21, 12, doi: 10.1186/s13058-018-1079-7 (2019).

7 Fievet, B. T. et al. Phosphoinositide binding and phosphorylation act sequentially in the activation mechanism of ezrin. J Cell Biol 164, 653–659, doi: 10.1083/jcb.200307032 jcb.200307032 [pii] (2004).

8 Niggli, V., Andreoli, C., Roy, C. & Mangeat, P. Identification of a phosphatidylinositol-4,5-bisphosphate-binding domain in the N-terminal region of ezrin. FEBS Lett 376, 172–176, doi: 10.1016/0014-5793(95)01270-1 (1995).

9 Roch, F. et al. Differential roles of PtdIns(4,5)P2 and phosphorylation in moesin activation during Drosophila development. J Cell Sci 123, 2058–2067, doi:123/12/2058 [pii] 10.1242/jcs.064550 (2010).

10 Roubinet, C. et al. Molecular networks linked by Moesin drive remodeling of the cell cortex during mitosis. J Cell Biol 195, 99–112, doi: 10.1083/jcb.201106048 (2011).

11 Tsukita, S. et al. ERM family members as molecular linkers between the cell surface glycoprotein CD44 and actin-based cytoskeletons. J Cell Biol 126, 391–401, doi: 10.1083/jcb.126.2.391 (1994).

12 Wu, K. L. et al. The NHE1 Na+/H+ exchanger recruits ezrin/radixin/moesin proteins to regulate Akt-dependent cell survival. J Biol Chem 279, 26280–26286, doi: 10.1074/jbc.M400814200 (2004).

13 Hoeflich, K. P. et al. Insights into a single rod-like helix in activated radixin required for membrane-cytoskeletal cross-linking. Biochemistry 42, 11634–11641, doi: 10.1021/bi0350497 (2003).

14 Turunen, O., Wahlstrom, T. & Vaheri, A. Ezrin has a COOH-terminal actin-binding site that is conserved in the ezrin protein family. J Cell Biol 126, 1445–1453 (1994).

15 Hamada, K., Shimizu, T., Matsui, T., Tsukita, S. & Hakoshima, T. Structural basis of the membrane-targeting and unmasking mechanisms of the radixin FERM domain. EMBO J 19, 4449–4462, doi: 10.1093/emboj/19.17.4449 (2000).

16 Coscoy, S. et al. Molecular analysis of microscopic ezrin dynamics by two-photon FRAP. Proc Natl Acad Sci U S A 99, 12813–12818, doi: 10.1073/pnas.192084599 192084599 [pii] (2002).

17 Pelaseyed, T. et al. Ezrin activation by LOK phosphorylation involves a PIP2-dependent wedge mechanism. Elife 6, doi: 10.7554/eLife.22759 (2017).

18 Schann, S., Bouvier, M. & Neuville, P. Technology combination to address GPCR allosteric modulator drug-discovery pitfalls. Drug Discov Today Technol 10, e261–267, doi: 10.1016/j.ddtec.2012.09.008 (2013).

19 Breton, B. et al. Multiplexing of multicolor bioluminescence resonance energy transfer. Biophys J 99, 4037–4046, doi: 10.1016/j.bpj.2010.10.025 (2010).

20 Avet, C. et al. Selectivity Landscape of 100 Therapeutically Relevant GPCR Profiled by an Effector Translocation-Based BRET Platform. 2020.2004.2020.052027, doi: 10.1101/2020.04.20.052027 %J bioRxiv (2020).

21 Benredjem, B. et al. Exploring use of unsupervised clustering to associate signaling profiles of GPCR ligands to clinical response. Nature communications 10, 4075, doi: 10.1038/s41467-019-11875-6 (2019).

22 Namkung, Y. et al. Functional selectivity profiling of the angiotensin II type 1 receptor using pathway-wide BRET signaling sensors. Science signaling 11, doi: 10.1126/scisignal.aat1631 (2018).

23 Schonegge, A. M. et al. Evolutionary action and structural basis of the allosteric switch controlling beta2AR functional selectivity. Nature communications 8, 2169, doi: 10.1038/s41467-017-02257-x (2017).

24 Namkung, Y. et al. Monitoring G protein-coupled receptor and beta-arrestin trafficking in live cells using enhanced bystander BRET. Nature communications 7, 12178, doi: 10.1038/ncomms12178 (2016).

25 Kondo, T. et al. ERM (ezrin/radixin/moesin)-based molecular mechanism of microvillar breakdown at an early stage of apoptosis. J Cell Biol 139, 749–758, doi: 10.1083/jcb.139.3.749 (1997).

26 Tachibana, K., Haghparast, S. M. & Miyake, J. Inhibition of cell adhesion by phosphorylated Ezrin/Radixin/Moesin. Cell Adh Migr 9, 502–512, doi: 10.1080/19336918.2015.1113366 (2015).

27 Liu, D. et al. Single-molecule detection of phosphorylation-induced plasticity changes during ezrin activation. FEBS Lett 581, 3563–3571, doi: 10.1016/j.febslet.2007.06.071 (2007).

28 Kobayashi, H., Picard, L. P., Schonegge, A. M. & Bouvier, M. Bioluminescence resonance energy transfer-based imaging of protein-protein interactions in living cells. Nature protocols 14, 1084–1107, doi: 10.1038/s41596-019-0129-7 (2019).

29 Carreno, S. et al. Moesin and its activating kinase Slik are required for cortical stability and microtubule organization in mitotic cells. J Cell Biol 180, 739–746, doi:jcb.200709161 [pii] 10.1083/jcb.200709161 (2008).

30 Machicoane, M. et al. SLK-dependent activation of ERMs controls LGN-NuMA localization and spindle orientation. J Cell Biol 205, 791–799, doi: 10.1083/jcb.201401049 (2014).

31 Canals, D., Roddy, P. & Hannun, Y. A. Protein phosphatase 1alpha mediates ceramide-induced ERM protein dephosphorylation: a novel mechanism independent of phosphatidylinositol 4, 5-biphosphate (PIP2) and myosin/ERM phosphatase. J Biol Chem 287, 10145–10155, doi: 10.1074/jbc.M111.306456 (2012).

32 Zhang, J. H., Chung, T. D. & Oldenburg, K. R. A Simple Statistical Parameter for Use in Evaluation and Validation of High Throughput Screening Assays. J Biomol Screen 4, 67–73, doi: 10.1177/108705719900400206 (1999).

33 Yang, H. S. & Hinds, P. W. Increased ezrin expression and activation by CDK5 coincident with acquisition of the senescent phenotype. Molecular cell 11, 1163–1176, doi: 10.1016/s1097-2765(03)00135-7 (2003).

34 Ren, L. & Khanna, C. Role of ezrin in osteosarcoma metastasis. Adv Exp Med Biol 804, 181–201, doi: 10.1007/978-3-319-04843-7_10 (2014).

35 Picard, L. P., Schonegge, A. M., Lohse, M. J. & Bouvier, M. Bioluminescence resonance energy transfer-based biosensors allow monitoring of ligand- and transducer-mediated GPCR conformational changes. Commun Biol 1, 106, doi: 10.1038/s42003-018-0101-z (2018).

