## Supplementary Figures for "Development of conformational BRET biosensors that monitor Ezrin, Radixin and Moesin activation in real-time"

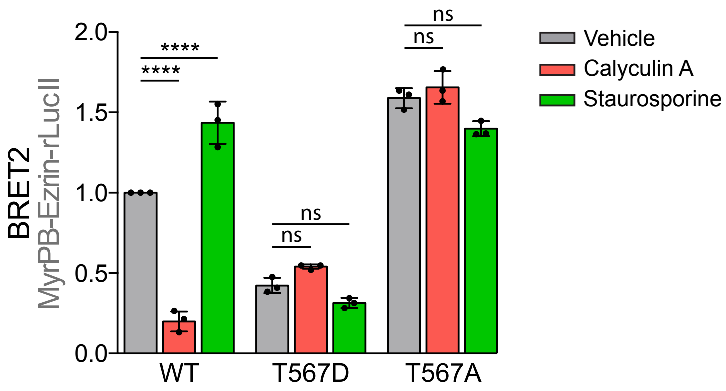


**Figure S1: MyrPB-Ezrin-rLucII mutants for the conserved regulatory residue are no longer sensitive to phosphatase/kinase inhibitors.**

HEK293T cells were co-transfected with rGFP-CAAX and a wild type (WT) or a constitutively active (T567D) or inactive (T567A) mutant of MyrPB-Ezrin-rLucII. Cells were then treated with either calyculin A or staurosporine. Graph represents the mean +/- s.d. of three independent experiments. Dots represent independent experiments. P values were calculated using one-way ANOVA, Tukey's multiple comparisons test with a single pooled variance. ****, $P < 0.0001$. ns, not significant.

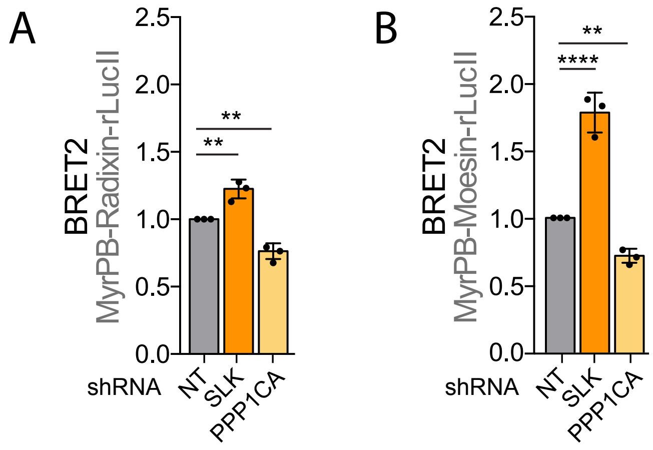


**Figure S2: Depletion of SLK and PPP1CA affects MyrPB-Radixin-rLucII and MyrPB-Moesin-rLucII opening measured by BRET.**

HEK293T cells knock-downed for SLK or PPP1CA were co-transfected with rGFP-CAAX and MyrPB-Radixin-rLucII (A) or MyrPB-Moesin-rLucII (B). Graphs represent the mean +/- s.d. of three independent experiments. Dots represent independent experiments. P values were calculated using one-way ANOVA, Tukey's multiple comparisons test with a single pooled variance. **, $P < 0.01$. ****, $P < 0.0001$.
